# Photosynthetic acclimation and sensitivity to short- and long-term environmental changes

**DOI:** 10.1101/2021.01.04.425174

**Authors:** Leonie Schönbeck, Charlotte Grossiord, Arthur Gessler, Jonas Gisler, Katrin Meusburger, Petra D’Odorico, Andreas Rigling, Yann Salmon, Benjamin D. Stocker, Roman Zweifel, Marcus Schaub

## Abstract

- The future climate will be characterized by an increase in frequency and duration of drought and warming that exacerbates atmospheric evaporative demand. How trees acclimate to long-term soil moisture changes and whether these long-term changes alter trees’ sensitivity to short-term (day to months) variations of vapor pressure deficit (VPD) and soil moisture is largely unknown.
- Leaf gas exchange measurements were performed within a long-term (17 years) irrigation experiment in a Scots pine-dominated forest in one of Switzerland’s driest areas on trees in naturally dry (control), irrigated, and‘irrigation-stop’ (after 11 years of irrigation) conditions.
- Seventeen years of irrigation increased photosynthesis (A) and stomatal conductance (g_s_) and reduced the g_s_ sensitivity to increasing VPD but not to soil drying. Following irrigation-stop, gas exchange did not decrease immediately, but after three years, had decreased significantly in irrigation-stop trees. Vc_max_ and J_max_ recovered after five years.
- These results suggest that long-term release of soil drought reduces the sensitivity to atmospheric evaporative demand and that atmospheric constraints may play an increasingly important role in combination with soil drought. In addition, they suggest that structural adjustments lead to an attenuation of initially strong leaf-level acclimation to strong multiple-year drought.

## Introduction

Many temperate ecosystems will experience an increase in frequency and intensity of both soil and atmospheric drought due to changing precipitation patterns and increasing temperatures that exacerbate atmospheric evaporative demand (higher vapor pressure deficit, VPD) (Simmons *et al.*, 2010; Willett *et al.*, 2014). To date, it is poorly understood how these combined stressors affect tree productivity, specifically how they affect the photosynthetic machinery of trees.

Knowledge of species-specific traits related to carbon assimilation and water consumption is crucial to assess and predict tree and forest functioning. Terrestrial biosphere models (TBMs) commonly use prescribed and temporally constant traits for a discrete set of plant functional types to specify photosynthetic capacity. The Farquhar, von Caemmerer and Berry (FvCB) model is widely used for mechanistically simulating photosynthesis (Farquhar *et al.*, 1980) as a function of photosynthetic capacities - Rubisco carboxylation (Vc_max_) and electron transport (J_max_). In addition, it relies on stomatal conductance (g_s_) and its sensitivity to the environment, which need to be estimated empirically or by modelling approaches funded in optimality principles for balancing carbon gains and water losses (Wang et al., 2020; Prentice et al., 2014; Medlyn et al., 2011; Wright & Westoby, 2003; Cowan & Farquhar, 1977). Common to most stomatal optimization models and the state-of-the-art in global TBMs is that Vc_max_ and J_max_ are assumed to be constants when expressed at a standard temperature and do not respond to varying soil moisture or VPD (Medlyn *et al.*, 2002; Egea *et al.*, 2011; De Kauwe *et al.*, 2013). Yet, empirical evidence in saplings and relative short-term experiments exists for acclimating responses in Vc_max_ and J_max_, e.g., during progressively drying soil conditions (Zhou *et al.*, 2014). However, the lack of long-term data on adult trees hampers our ability to estimate drought impacts and understand the susceptibility and capacity for acclimation of plant carbon assimilation.

Short-term acclimation occurs when trees adjust their physiology to overcome slowly increasing stresses (Kozlowski and Pallardy, 2002; Marchin *et al.*, 2016; Grossiord *et al.*, 2018). Stomata regulate the balance between carbon intake and water loss. They will close (g_s_ decreases) when trees experience soil or atmospheric drought, which will reduce transpiration (E), CO_2_ diffusion, and CO_2_ concentration inside the leaf (Ci). Lower Ci, in turn, causes reduced leaf-level photosynthetic activity (A). Drought could also lead to a decrease in Rubisco activity, down-regulating the activation state of the enzyme, leading to a reduction in Rubisco content and/or soluble protein content (Parry, 2002). While stomatal closure, and thus stomatal limitation, is purely affecting diffusion of CO_2_ and water vapor, other processes, summarized as non-stomatal limitation of photosynthesis, affect the diffusion of CO_2_ through the mesophyll, and the photo- and biochemistry of photosynthesis. For example, the downregulation of Rubisco can be considered as a non-stomatal but biochemical limitation of the photosynthetic capacity during drought, leading to reduced Vc_max_ and J_max_ (Kanechi *et al.*, 1996; Wilson *et al.*,2000; Castrillo *et al.*, 2001; Parry, 2002; Tezara, 2002; Zhou *et al.*, 2014). Moreover, mesophyll conductance (g_m_) can exert a diffusional but non-stomatal limitation on photosynthesis, and, depending on the species, increases, decreases or doesn’t change at all in response to drought (Hommel *et al.*, 2014). Several studies found that deciduous tree species during drought mainly showed stomatal limitation of A (Wilson *et al.*, 2000; Flexas *et al.*, 2004; Keenan *et al.*, 2010), while other studies found that the effect size of the two limitations is strongly dependent on tree species, habitat, and duration of drought (Zhou *et al.*, 2013, 2014; Salmon *et al.*, 2020).

Stomatal and biochemical acclimation could be achieved on a timescale from minutes to weeks, whereas structural acclimation, such as adjustment of root-to-leaf ratio or leaf-to-sapwood area, could take multiple years (Sultan, 2000; Poyatos *et al.*, 2007; Martínez-Vilalta *et al.*, 2009). Long-term structural changes can alter the sensitivity of the stomata and photosynthetic apparatus to short-term fluctuations in the environment (e.g. Gessler *et al.*, 2017). For example, a high ratio of leaf-to-sapwood area resulting from acclimation to high soil water availability allows for similar C assimilation with increased water loss (Zweifel *et al.*, 2020) but may pose a hydraulic risk during sudden heatwaves with strong atmospheric demand, causing the stomata to close rapidly. Moreover, a reduced ratio of root-to-leaf area might expose the tree to greater risk during extreme and enduring soil drought. In contrast, trees acclimated to low soil water content perform a more conservative water use. Their lower leaf area reduces total tree water loss while enabling leaves to maintain their function with normal g_s_ and A (Pataki *et al.*, 1998; Kelly *et al.*, 2016). They might capitalize on sudden increases in soil water, while their posture with smaller leaf area will probably not react as strongly on fluctuations in evaporative demand. Also, Vc_max_ and J_max_ might be prone to acclimation where acclimation to aridity might lead to significantly more protein allocated to Rubisco in leaves (Pankovic *et al.*, 1999; Wright *et al.*,2003; Prentice *et al.*, 2014; Wang *et al.*, 2017). Still, how exactly photosynthetic acclimation to changing soil moisture proceeds, and how acclimation affects the sensitivity of g_s_ to short-term environmental fluctuations is unknown.

There is an increasing need to characterize tree physiological sensitivity and acclimation to atmospheric and soil drought (Grossiord *et al.*, 2020). Measuring leaf-level photosynthetic capacity and sensitivity to environmental cues is time-consuming but indispensable for answering to which extend trees respond to drought over the long *vs.* the short term. Although leaf-level gas exchange measurements have been conducted on multiple species, no study has yet attempted to decipher how long-term exposure to soil moisture change, and subsequent adjustments to novel conditions, could alter the sensitivity of photosynthetic properties to environmental variability. In a Scots pine-dominated forest in one of Switzerland’s driest areas, we conducted leaf-level gas exchange measurements in a long-term irrigation experiment covering multiple years. We tested (1) how photosynthetic properties (i.e., A, g_s_, J_max_, and Vc_max_) acclimate in response to long-term (17 years) artificial change in soil moisture (naturally drought-exposed control trees *vs.* irrigated trees), (2) how acclimation to long-term changes in soil moisture impacts the sensitivity of photosynthetic properties to short-term VPD and soil moisture variation, and (3) how fast photosynthetic properties of trees recover when drought follows a long-term acclimation to high soil moisture (irrigation stopped after 11 years). We hypothesized that (1) acclimation to long-term irrigation had led to similar A, g_s_, Vc_max_ and J_max_ and lower intrinsic water use efficiency (WUE_i_) compared to control trees, due to structural acclimation; (2) irrigated trees will show stronger sensitivity to atmospheric drivers due to their sizeable water-consuming crown, while control trees will react stronger to soil moisture fluctuations; (3) trees released from the irrigation and exposed to sudden drought (irrigation-stop) will strongly reduce their photosynthetic properties in the first year, followed by structural adjustments (e.g., lower crown leaf area and water-conducting area) that allow for a recovery of leaf-level gas exchange.

## Materials and Methods

### Site and experimental design

A 17-year irrigation experiment was conducted in the Pfynwald forest (46°18N, 7°36’ E, 615 m a.s.l.), the largest Scots pine *(Pinus sylvestris* L.) dominated forest in Switzerland, located in the dry inner-Alpine valley of the river Rhone, close to the dry edge of the natural distribution of Scots pine (Critchfield and Little, 1966). The Pfynwald is a 100-year-old naturally regenerated forest, but past forest practices may have favored Scots pine regeneration over other species such as *Quercus pubescens* (Weber *et al.*, 2008; Gimmi *et al.*, 2010; Rigling *et al.*,2013). Climatic conditions are characterized by a mean annual temperature of 10.1 °C and a yearly precipitation sum of approximately 600 mm. Scots pine forests in this region are regularly subjected to drought- and heat-induced mortality (Bigler *et al.*, 2006; Allen *et al.*,2010; Rigling *et al.*, 2013). The average tree age is approximately 100 years, and the forest has a mean canopy height of 10.8 m, a stand density of 730 stems ha^-1^, and a basal area of 27.3 m^2^ ha^-1^ (Dobbertin *et al.*, 2010). The soil is a calcaric regosol (FAO classification) characterized by very low water retention and high vertical drainage (Brunner *et al.*, 2009).

The experimental site (1.2 ha; 800 trees) is divided into eight plots of 25 m x 40 m each, separated by a 5 m buffer zone. The irrigation of ~600 mm/year is applied at night on four plots between April and October, from 2003 onwards, with 1 m high sprinklers using water from a nearby channel running parallel to the experimental plot, fed by the Rhone River. Nutrient input through irrigation was proven to be minor (Thimonier *et al.*, 2005, 2010). In 2014, irrigation was stopped in the upper third of the irrigated plots, resulting in three categories: controls (non-irrigated) representing the natural dry condition; irrigation resulting in a release of soil drought; irrigation-stop exposing trees that were acclimated to well-watered conditions for 11 years to drought (Fig. 1). In 2015, nine scaffolds were installed in the forest, three per treatment, to enable easier access to tree crowns for sampling and *in situ* measurements. The volumetric soil water content (VWC) was monitored hourly in one control and one irrigated plot until 2014, using time domain reflectometry (Tektronix 1502B cable tester, Beaverton, OR, USA), at a soil depth of 10, 40, and 60 cm at four different locations per plot. In 2014, all soil moisture sensors were replaced with Decagon 10-HS sensors (Decagon Devices, Inc., Pullman, WA, USA). They were installed in six different plots (two irrigated, two control, and two irrigation-stop plots), at 10 and 80 cm depth. Air temperature, relative humidity (Sensirion SHT-21, Sensirion AG, Switzerland), and precipitation (Tipping Bucket Rain Gauge, R.M. Young, MI USA) were measured on-site.

**Figure 1:**
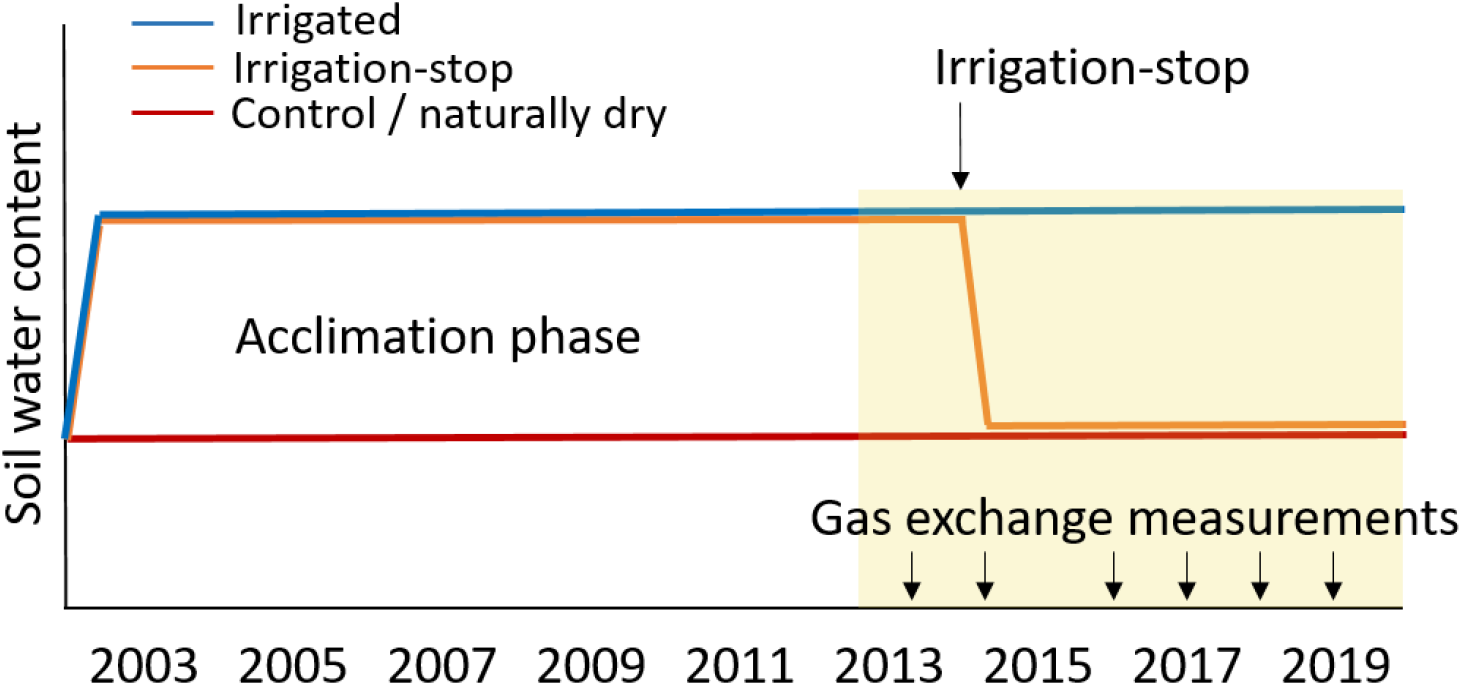
Timeline of treatments and measurements from 2003 to 2019 in a long-term irrigation experiment. Irrigation was stopped in 2014 in 1/3 of the irrigated plots. Gas exchange measurements took place three times a year, from 2013 to 2019, except in 2015. In 2016, only control and irrigated trees were measured.

### Gas exchange measurements

In 2013, 2014, and 2016-2019, leaf gas exchange measurements were carried out in the form of A/C_i_ measurements. In 2013 and 2014, hunting seats were used on specific trees to access fully sun-exposed branches of the outer upper crown for leaf-level gas exchange measurements. In 2016, gas exchange was measured on branches cut off from sun-exposed parts in the upper half of the canopy, cut again underwater, and kept in a bucket of water. In 2017-2019, measurements were conducted from the top of the nine scaffolds and on three trees per scaffold. Here, the top of the canopy was reached, and sun-exposed needles were selected for measurements. Although measurements were taken from slightly different canopy depths but always from sun-exposed branches from the outer crown, we assumed no strong gradients in light, VPD, or other environmental conditions within the sparse canopy of the trees at our site. In a Scots pine stand with a comparable structure, no intra-canopy gradients in gas exchange were observed (Brandes *et al.*, 2006). Thus, we contend that the branches selected from the outer crown were comparable and representative of the whole canopy. Until 2019, measurements were carried out with two LiCor LI-6400 systems (LiCor Inc., Lincoln, Nebraska, USA). The instruments were replaced by the LI-6800 system (LiCor Inc.) in 2019. A/C_i_ measurements were taken once in spring (May/June), summer (July/August), and autumn (October) of each year. Additional point measurements at 400 ppm CO_2_ were taken in 2013, 2014, and in the summer of 2017, which were included in the analyses regarding A and g_s_. Five to 10 one-year-old (i.e., previous year) needles were enclosed clipped in the cuvette gasket, ordered in a flat plane without overlapping each other. The temperature inside the cuvette was set close to the outside midday temperature. With the LI-6800, VPD was set to 1.5 kPa, while in the LI-6400, humidity regulation was not a built-in function, and RH was maintained between 60-70%. In most cases, this led to a decrease in VPD in the cuvette compared to outside conditions (Supporting information Fig. S1). Nonetheless, the complete dataset comprises a range of VPD values between 0.3 and 3 kPa. The actual conditions the needles experienced, i.e. the cuvette VPD, was used as a covariate for statistical analyses. Photosynthetically active radiation (PAR) was kept at saturation point of 1000-1200 μmol m^−2^ s^−1^ (Palmroth and Hari, 2001).

Photosynthetic activity was measured at CO_2_ concentrations in the sequence steps of 400, 300, 200, 100, 50, 0, 400, 600, 800, 1200, and 1800 ppm. After each measurement, the part of the needles enclosed in the cuvette was harvested, and the projected leaf area (Serrano *et al.*, 1997; Renninger *et al.*, 2015) was measured using a flatbed scanner and analyzed using Pixstat (Pixstat v1.3.0.0, Schleppi 2018). The projected leaf area of the measured foliage was used to correct the recorded gas exchange values.

### A/C_i_ Curve fitting

A/C_i_ curves were fitted using the Farquhar, von Caemmerer & Berry model for photosynthesis, described in Sharkey et al. (2007) and computed in the ‘plantecophys’ package (Duursma, 2015). Before fitting, the data was cleaned based on visual determination to get rid of unreasonable numbers due to measurement artifacts, following these criteria (Gu *et al.*, 2010):

- 0 ppm < C_i_ < 2000 ppm
- 0 mol/m^2^/s < g_s_ < 1.5 mol/m^2^/s
- -5 < A < 20 mmol/m^2^/s
- Each A/C_i_ curve must have reached a C_i_ of 600 ppm to ensure a saturating plateau
- A/C_i_ curve must have more than 5 points after the previous selection.

The ‘plantecophys’ package’s default method was used if possible; all other fits were done with the binomial method. The default assumption of infinite mesophyll conductance (g_m_) was used. As a result, ‘apparent’ Vc_max_ and J_max_ are computed, and changes in apparent Vc_max_ and J_max_ reflect changes in both biochemical limitations as well as mesophyll conductance. The model used a temperature correction to fit all curves to 25°C. Transition point, i.e. the C_i_ where the transition takes place from Rubisco limited photosynthesis to RuBP regeneration/electron transport limitation, was estimated by the model, as well as day respiration (R_d_), photorespiratory compensation point (Γ*) and the Michaelis-Menten Coefficient (Km, Pa) (Supplementary data Table S1). After fitting the curves, non-fitting curves were eliminated following the following criteria based on validated values in the literature (von Caemmerer and Farquhar, 1981; Wullschleger, 1993; Gu *et al.*, 2010) and by visual determination of extreme outliers:

- 0 ppm < Transition point (T_p_) < 1600 ppm
- 3 *μ*mol/m^2^/s < J_max_ < 150 *μ*mol/m^2^/s
- 1 *μ*mol/m^2^/s < Vc_max_ < 95 *μ*mol/m^2^/s
- Root mean squared error (measure of accuracy) < 10

After cleaning, 213 out of 312 measured curves were considered in the analyses.

### Statistical analysis

To test for general, long-term, treatment differences in A, g_s_, E, intrinsic water use efficiency (WUE_i_, A/g_s_), Vc_max_, and J_max_, a linear mixed effect model with the control and irrigated treatment as fixed and tree individual nested in year as a random factor was used. The mixed effect models were fitted using the ‘lmerTest’ package in R (Kuznetsova et al., 2017). The year was also included as a fixed factor but never interacted with the treatment. It was thus chosen to focus on the general treatment effect.

Linear mixed effect models were also used to test whether the long-term manipulation of soil water changed the sensitivity of leaf g_s_ to short-term environmental variations. Only trees from control and irrigated treatments were used for this analysis. The widely described relationship between g_s_ and VPD was used as a basis for the g_s_ model. g_s_ is described using (Oren *et al.*, 1999):

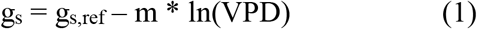

Where g_s_ is the stomatal conductance at any level of VPD, g_s,ref_ is the reference stomatal conductance at a VPD of 1 kPa, and m is the sensitivity of g_s_ to VPD. We then extended this model with soil volumetric water content (2-degree polynomial) and treatment as fixed factors, including all interactions, and tree nested in year as a random factor. Model selection was then made according to the lowest Akaike information criterion (AIC). If needed, variables were log- or square-root-transformed to meet the normal distribution of the residuals. For visualization, three soil VWC bins were created, splitting the data up in soil VWC of 25-40%, 41-55%, and 56-70%. Modeled data were simulated using the ‘arm’ package (Gelman *et al.*,2020).

Acclimation of all photosynthetic parameters previously described over time in the ‘irrigation-stop’ plots was analyzed by testing for a difference between irrigated and the irrigation-stop in a mixed effect model. Treatment and month were fixed factors, and tree nested in date was treated as a random factor. Using the ‘multcomp’ package in R (Hothorn *et al.*, 2019), pairwise comparisons per date were visualized.

## Results

### Climate

Volumetric soil water content at 10 and 80 cm depth was significantly higher in the irrigated plots than in the control and irrigation-stop plots during the summer months when the irrigation was activated (Fig. 2). From late autumn until spring, soil water content was comparable over all treatments because the irrigation treatment was switched off. Natural precipitation events (and failures in the irrigation system) during the summer months occasionally reduced the treatment differences across soil depths. Mean daily VPD ranged from close to 0 in the winter months to a maximum of 2.5 kPa in the dry and hot summer months.

**Figure 2:**
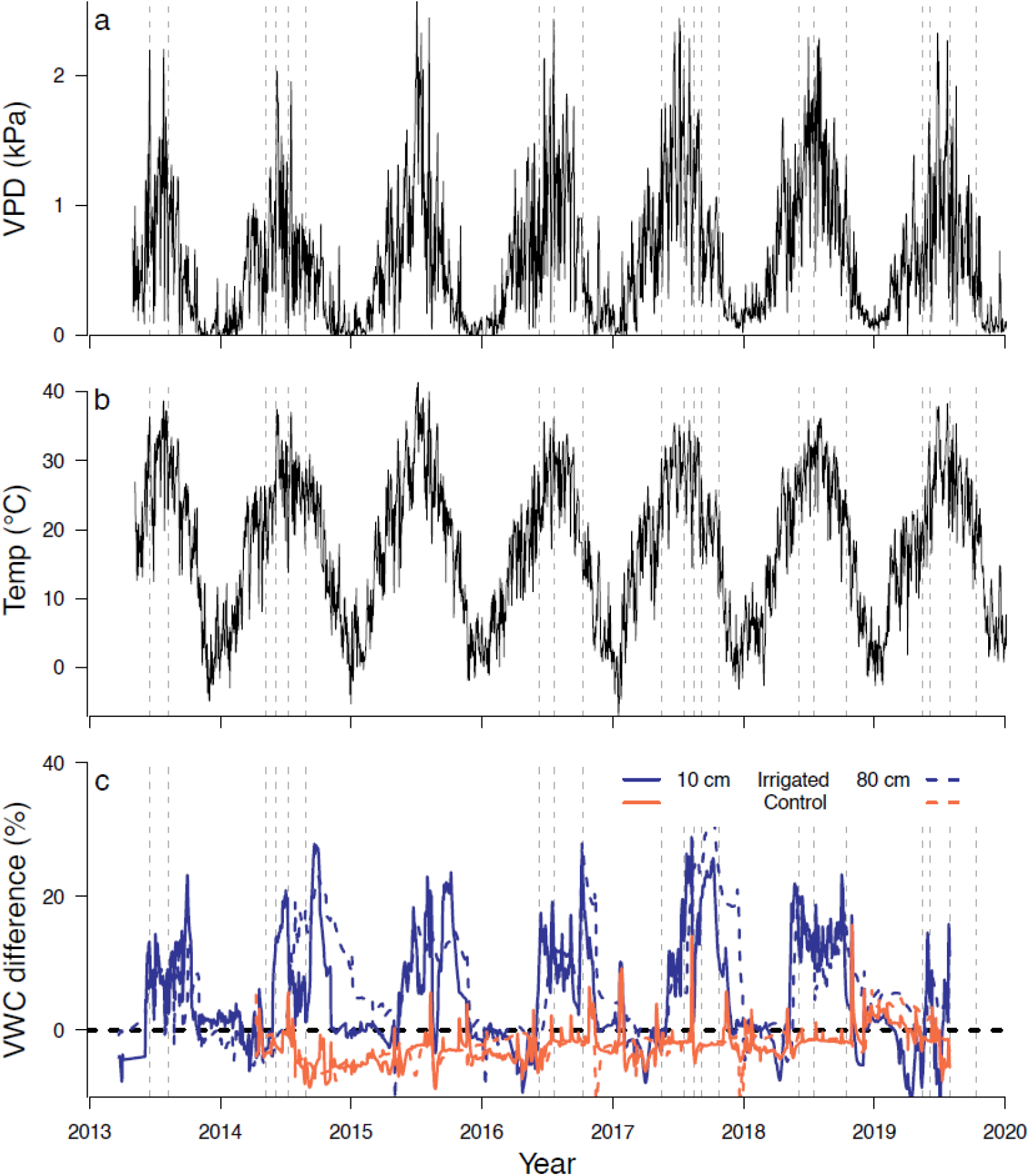
(a) Daily means of vapor pressure deficit (VPD), (b) maximum daily temperatures, and (c) differences in daily volumetric soil water content (VWC) between control and irrigated (blue), and control and irrigation-stop plots (orange). Solid lines show soil VWC at 10 cm depth and dashed lines at 80 cm depth. Grey vertical dashed lines indicate the dates when gas exchange measurement campaigns were conducted.

### Acclimation to long-term environmental conditions

Irrigated trees showed in general higher g_s_ (60% increase, p < 0.001), A (34% increase, p < 0.001) and E (60% increase, p < 0.001) than control trees after the 11-year acclimation period (Fig. 3, Table S2). WUE_i_ was only slightly lower in irrigated than in control trees (6% difference, p=0.04) and no significant treatment difference was found for Vc_max_, (C = 44.4 ± 2.14 *vs.* 43.9 ± 1.7) and J_max_ (C = 69.4 ± 2.9 *vs.* I = 75.8 ± 2.5), while the ratio between the two ^(J^max^/Vc^max^)^was marginally higher in irrigated trees compared to control trees (C = 1.6 ± 0.05 *vs*. I = 1.8 ± 0.06) (Fig. 3, Table S2).

**Figure 3:**
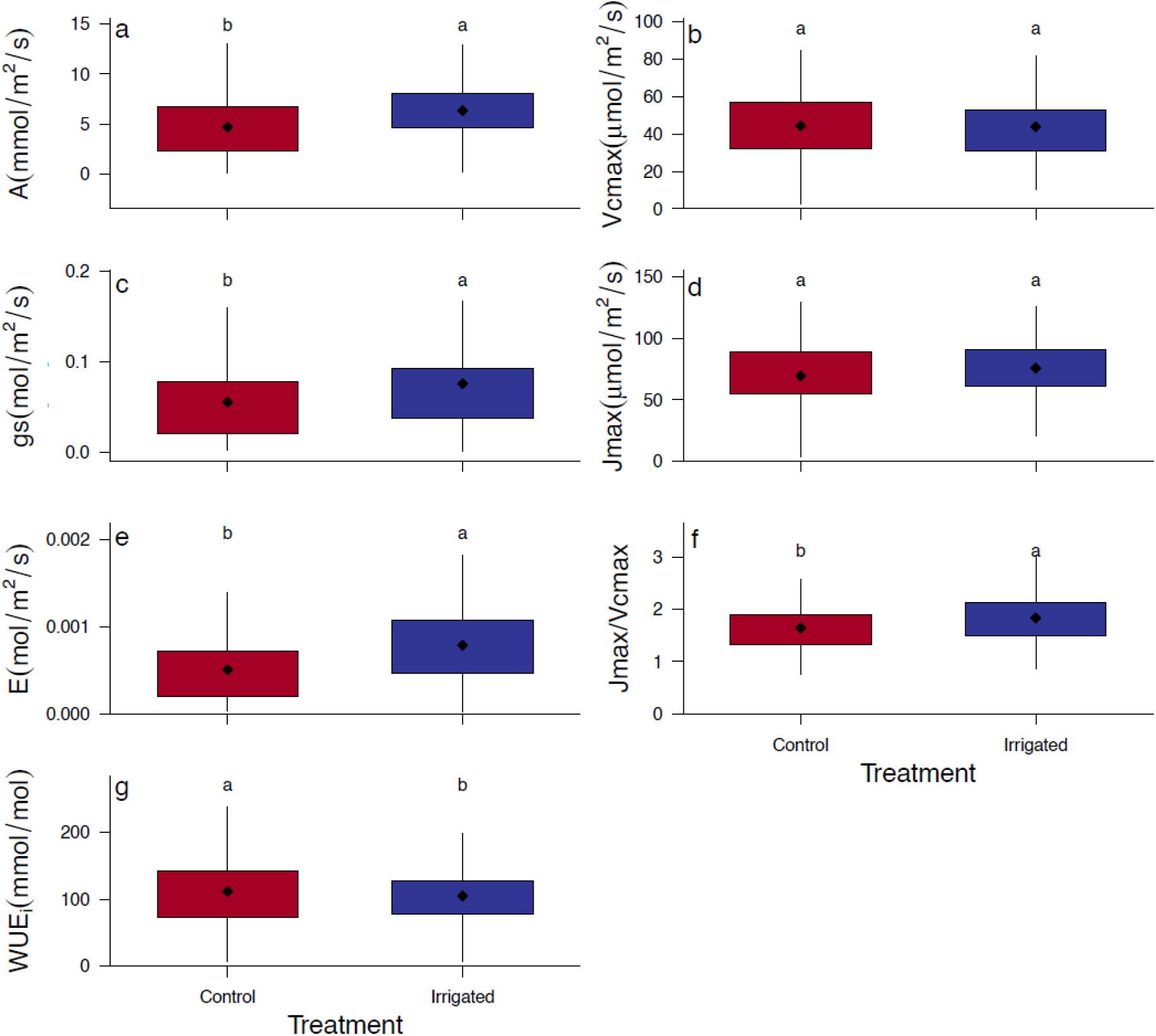
Treatment differences for photosynthesis (A), stomatal conductance (g_s_), transpiration (E), intrinsic water use efficiency (WUE_i_), Vc_max_, and J_max_ and the ratio between the two as an average of all measurements between 2013 and 2019 in control / natural dry (red) and irrigated (blue) trees. Symbols show the mean; boxes and whiskers the interquartile ± 1.5 * interquartile range. Different letters above the boxplots indicate significant group differences according to linear mixed effect models (p < 0.05).

### Sensitivity of g_s_ to short-term environmental changes

A strong logarithmic relationship was found between g_s_ and VPD (Fig. 4a). The long-term acclimation of the trees affected g_s_’ sensitivity (i.e., m, the slope of the curve) to short-term fluctuations in VPD. With increasing VPD, control trees reduced g_s_ faster than irrigated trees (Fig. 4a; Table S3, S4). This treatment difference was most pronounced in low and intermediate soil VWC (25 - 55% VWC) (Supporting information Fig. S2). Soil drying below 55% VWC resulted in a decrease of g_s,ref_ for both treatments (Fig. 4b).

**Figure 4:**
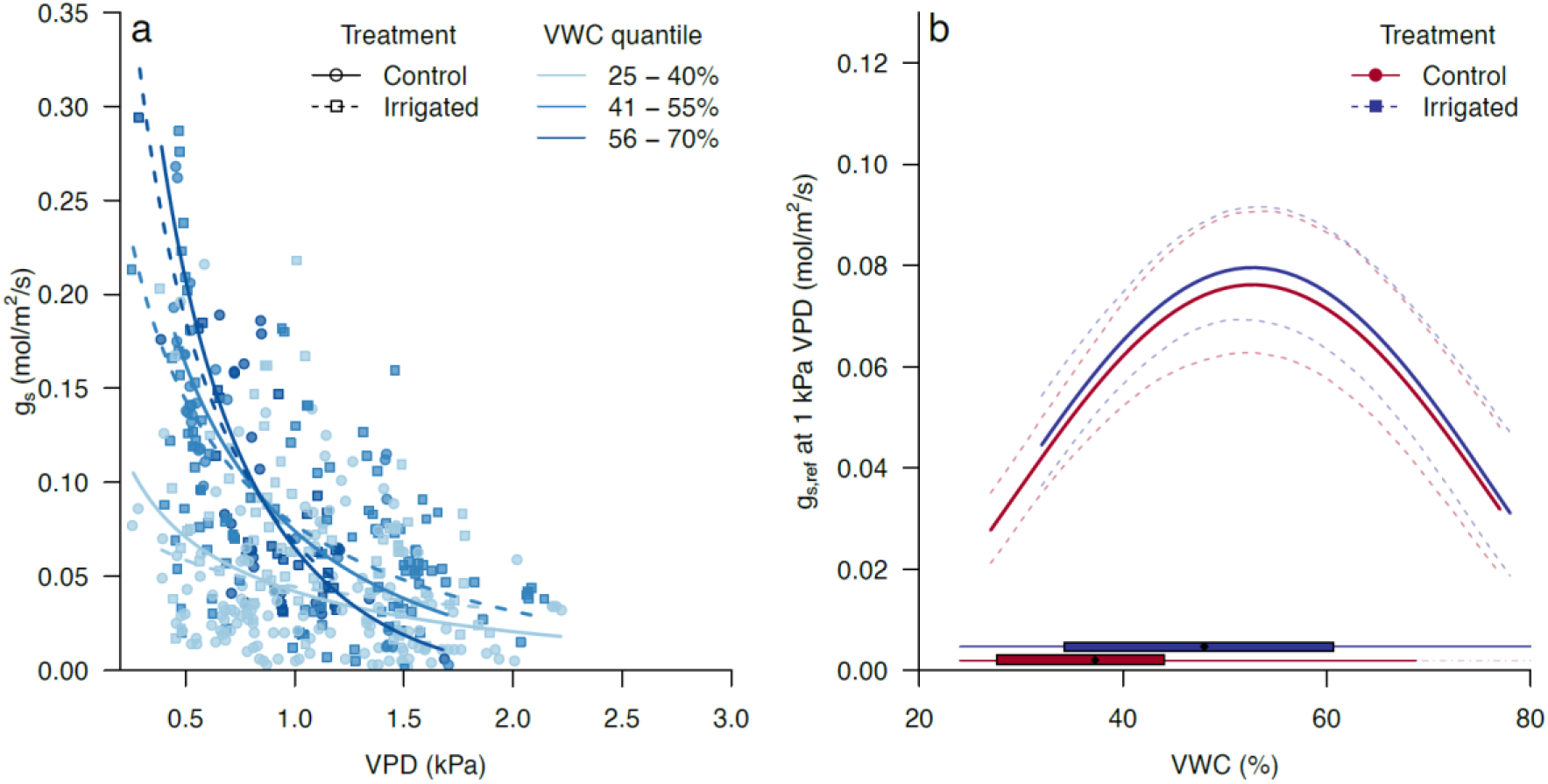
a) Stomatal conductance (g_s_) *vs.* vapor pressure deficit (VPD). Colors show fitted model regressions according to the linear mixed effect model, per volumetric soil water content (VWC) quantile. Line type distinguishes control and irrigated treatments. b) g_s,ref_ (g_s_ at 1 kPa VPD) as a function of soil VWC per treatment according to the linear mixed effect model. Colors indicate control and irrigated treatments. Dashed lines show the 95% credibility intervals of the mixed effect model. The range distribution of soil VWC in control and irrigated plots is shown on the bottom of the graph for reference.

### Acclimation over time after irrigation-stop

During 2014, the year of the irrigation-stop, assimilation did not decline significantly in the irrigation-stop trees compared to irrigated trees (Fig. 5, Supporting information Fig. S3). In spring 2014, A, Vc_max_ and J_max_ were even higher in irrigation-stop trees than in irrigated trees. From 2017 onwards, A, g_s_, and E were significantly lower in irrigation-stop than irrigated trees (Fig. 5) and had reduced significantly compared to 2014 (Supporting information Fig. S3; Supporting information Table S2). Water use efficiency showed a steady increase from 2017 to 2019; both compared to irrigated trees and in absolute terms (Fig. 5, Supporting information Fig. S3). Vc_max_ and J_max_ dropped from being higher than irrigated trees in 2014 to lower or comparable in 2017. Vc_max_ recovered to original levels in 2018 and 2019. The ratio between J_max_ and Vc_max_ decreased between 2014 and 2019 steadily but not significantly.

**Figure 5.**
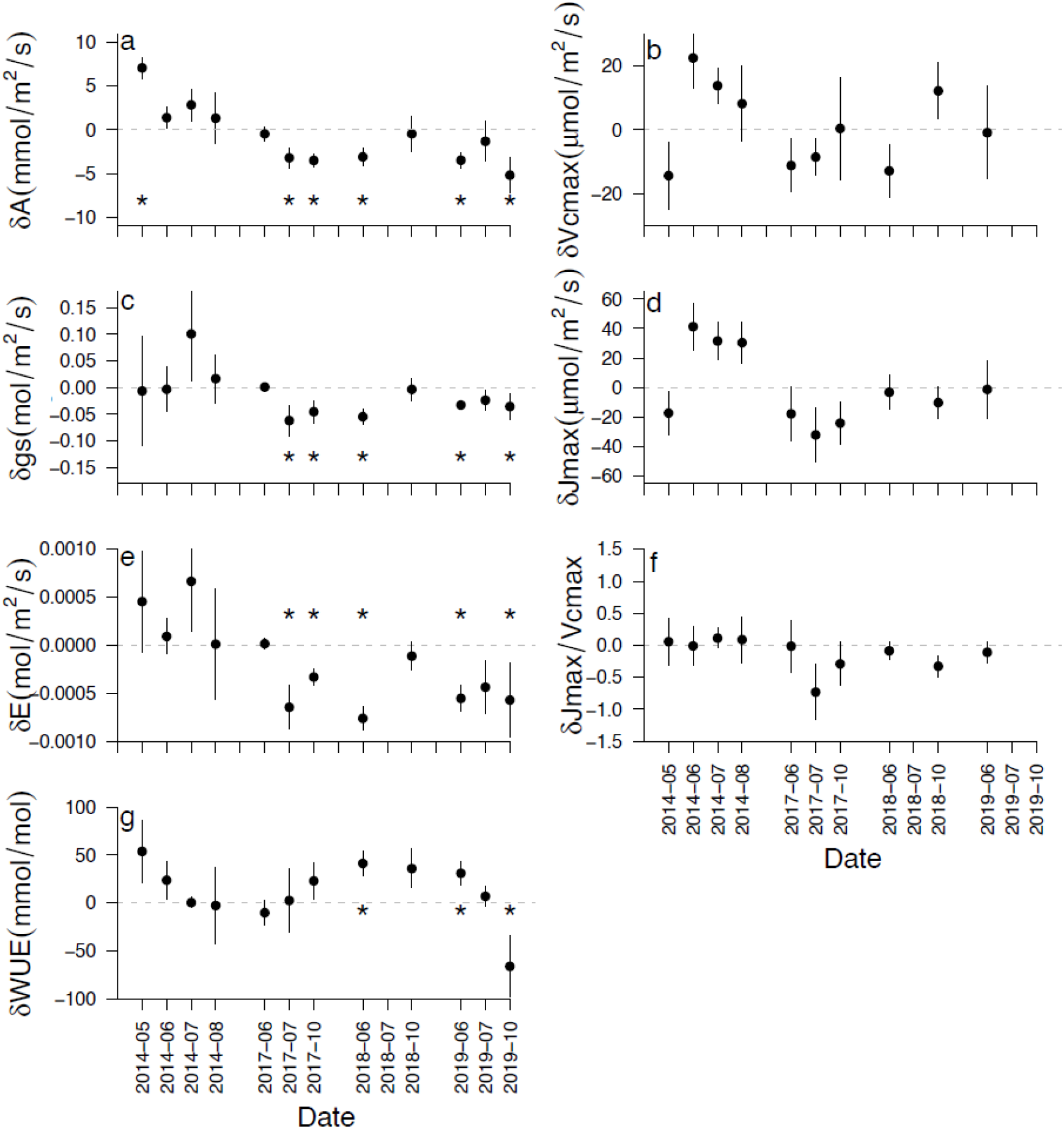
Differences between ‘irrigation-stop’ and irrigated trees at four measuring campaigns in 2014 and three in each year of 2017, 2018 and 2019. Plots show photosynthesis (a), stomatal conductance (g_s_) (c), transpiration (E) (e), intrinsic water use efficiency (WUE) (g), Vc_max_ (b) and J_max_ (d), and the ratio between the two (f). Symbols indicate the difference of the two means (irrigation-stop minus irrigated) and error bars the SE of the mean differences. Asterisks indicate significant differences between irrigated and irrigation-stop trees (p < 0.05). Data is missing in 2018-07 due to technical problems resulting in a low number of tree replicates and absence of measurements on irrigation-stop trees. A/C_i_ parameters are missing in July and October 2019, when only point measurements were taken.

## Discussion

### Long-term acclimation to environmental conditions

Our hypothesis that irrigated trees had lower water use efficiency (WUE_i_), and similar photosynthesis (A), stomatal conductance (g_s_), evaporation (E), Vc_max_ and J_max_ was confirmed for Vc_max_ and J_max_, and very weakly for WUE_i_, while A, g_s_, and E were even higher in irrigated than control trees. Since the start of irrigation in 2003, irrigated trees have increased total leaf area compared to the control trees (Schönbeck *et al.*, 2018). The higher assimilation rates together with structural acclimation suggest an even higher C turnover than expected in irrigated compared to control trees. The differences in gas exchange parameters are relatively small, even though they are statistically significant, which is probably caused by these structural adjustments in the canopy, which reduces the need for large assimilation increases and limits the variation in WUE_i_ between control and irrigated trees (Cinnirella *et al.*, 2002). A smaller crown (lower total leaf area) reduces the total water loss and reduces the need for leaf-level water-conserving strategies such as increasing WUE_i_ (Cinnirella *et al.*, 2002; McDowell *et al.*,2002). The fact that A and g_s_ were higher but Vc_max_ and J_max_ were similar in irrigated compared to control trees suggests a strong stomatal control of photosynthesis (Zhou *et al.*, 2013), with a smaller role for, and potential acclimation of, Rubisco activity (Parry, 2002), electron transport capacity (Epron and Dreyer, 1992) or mesophyll conductance (Egea *et al.*, 2011). This speculation is corroborated by the fact that nitrogen concentration in the leaves, an essential component of Rubisco, did not change with irrigation (Schönbeck *et al.*, 2018).

### Short-term sensitivity as affected by long-term acclimation to artificial change

We expected a higher sensitivity of g_s_ to VPD and soil VWC fluctuations in irrigated trees compared to control trees. Instead, control trees showed a higher sensitivity of g_s_ to increasing VPD. These results suggest that atmospheric constraints may play a more critical and increasingly important role in trees exposed to soil drought than trees growing in wetter soil conditions. This is in contradiction with a study by Novick *et al.* (2016), who predicted a smaller role for VPD limitation in soil moisture limited biomes. However, it is important to note that in our study, no comparison is made between biomes but between trees that have been exposed to a number of severe droughts in the last two decades, and trees that have been released from these drought episodes. Control trees constantly operate at significantly lower soil VWC than irrigated trees (37%*vs.* 48% resp., Fig. 4b). Although structural acclimation - i.e. smaller leaf area compared to irrigated trees - should reduce whole-tree transpiration and thus water demand of the control tree compared to irrigated trees, needle water potentials measured in 2016 show that control trees do experience lower water potentials and hence drought stress (Schönbeck *et al.*, 2018). *Pinus sylvestris* is an isohydric species, indicating a strong control of stomatal conductance with increasing VPD (Meinzer *et al.*, 2009; Martínez-Sancho *et al.*, 2017). In analyses of whole-tree sap flow, Grossiord *et al.* (2018) found similar sensitivity of sap flow in control and irrigated trees to VPD. However, they defined sensitivity only by the maximum sap flux density at optimal VPD, which is more comparable to g_s,ref_. We do agree that this influences the sensitivity curve. Still, we think that the sensitivity parameter *m*, which is significantly different between treatments in our study does indicate a higher sensitivity of g_s_ to VPD in control trees. It should be noted that extrapolation from the needle-level to the crown is very complex, and we do state that in our study, the highest strength lies in determining leaf-level photosynthetic characteristics. The sensitivity of g_s_ to soil VWC changes is well reported in other studies that show that soil drying reduces g_s,ref_ (Schäfer, 2011). For control trees, this means that they can keep their physiological potential to exploit ‘windows of opportunities’ - i.e. times when water is available in higher amounts. Such rapid responses to precipitation were demonstrated by Joseph et al. (2020), who found a strong increase in carbon allocation to belowground tissues in the control plots of the same forest system after a precipitation event.

### Acclimation to sudden long-term changes in precipitation

Against our expectation, the stop of irrigation did not reduce any leaf-level gas exchange parameter in the first year, 2014 in relation to the irrigated trees (Fig. 5), despite rapid reductions in the soil available water (Fig. 2). Instead, irrigation-stop trees kept similar g_s_, E, and A as irrigated trees. A slight reduction in A and E was observed between May and August 2014. It should be noted that the treatment differences in soil VWC became apparent only in June 2014. It is thus logical to expect no treatment differences in May 2014 yet. Irrigation-stop trees had even higher A than irrigated trees in May 2014. Over the summer months, the continued high evaporation rates and stomatal conductance could have translated into even lower soil VWC in the irrigation-stop plots than in control plots from July 2014 onwards. These findings do not fully correspond to the results of a study on sap flow and tree water deficit by Zweifel *et al.*(2020). They found a gradual decrease of whole tree sap flow rates over the season and even lower values than control trees from June onwards. Whole-tree sap flow measurements are a result of transpiration of the entire crown. In our study, we always measured needles that emerged in the previous year. Thus, a significant physiological and morphological difference between newly emerged needles and the older cohorts was created during the year 2014. Indeed, Zweifel *et al.* (2020) show that needle length decreased to levels below those of control trees, significantly reducing the total leaf area of the tree. Several studies show large differences between needle cohorts in coniferous species (Jach and Ceulemans, 2000; Robakowski and Bielinis, 2017), and as the year advances towards the end of the growing season, current year needles become increasingly important for tree productivity (Jensen *et al.*, 2015). The steep reduction in the newly formed leaf area could have had significant impacts on the whole crown transpiration, while the older needle cohorts did not react to the changes in soil VWC yet. Similar findings were reported

Three years after the halt of irrigation, A, g_s_, and E had dropped below the levels of irrigated trees and remained lower for most measuring dates until the end of 2019. Water use efficiency showed a gradual increase over time until 2018 and dropped again in 2019. The slow process of increasing WUE_i_ is surprising; however, it is a sign for a conservative but very plastic acclimation strategy of pine (Zweifel and Sterck, 2018). WUE_i_ is highly variable over the season, but an average increase of WUE_i_ relative to irrigated trees was expected in the first few needle cohorts. Instead, WUE_i_ in irrigation-stop trees was higher than in irrigated trees for the first time in June 2018 (measured on the needle cohort emerging in 2017). It appears that the characteristics of newly built tree structures are programmed not only by current conditions but also by a certain ecological memory effect, which limits the range of adjustments of trees to environmental changes (Anderegg *et al.*, 2018; Zweifel *et al.*, 2020).

Apparent J_max_ and Vc_max_ had also dropped (Supporting information Fig. S3) to slightly lower levels compared to irrigated trees (Fig. 5) but seemed to have recovered in 2018 again. While it was expected for A, g_s_, and E, the effects of drought on apparent Vc_max_ and apparent J_max_ are far less understood. These results indicate that both stomatal/diffusional limitations and biochemical limitations inhibited photosynthesis in 2017. Decreases of J_max_ have been observed for plant species across many ecosystems (Nogués and Baker, 2000; Ogaya and Peñuelas, 2003; Pezner *et al.*, 2020), suggesting that many species experience a downregulation of electron transport in response to drought. Interestingly, apparent Vc_max_ and, in a lesser amount, apparent J_max_ show a recovery over the years 2017-2019. Such a recovery was also found in some other species (Pankovic *et al.*, 1999; Damour *et al.*, 2009; Zhou *et al.*, 2016) and could be a result of structural acclimation to drought during the three years of irrigation-stop, allowing for higher ‘per-leaf-area’ gas exchange rates (McDowell*et al.*, 2002; Schönbeck *et al.*, 2018). For Scots pine, a full adjustment of the crown would take approximately 3-5 years, corresponding to the total number of needle cohorts from a tree crown (Zweifel *et al.*, 2020). This shows that the photosynthetic capacity on a biochemical level has recovered so that ‘windows of opportunity’ can be fully optimized, while on average, leaf-level A, E, and g_s_ remain low - i.e. comparable to control trees.

This study was carried out over seven years, which creates a highly needed long-term perspective of leaf acclimation to changing environmental conditions. It highlights the importance of understanding leaf structure and biochemical composition to better extrapolate whole-tree water and carbon dynamics. Compared to other spatially large-scale studies, the results show that more knowledge is needed on between-needle-cohort differences in structure and function to translate these results to whole-crown or whole-tree hydraulics and carbon dynamics. Nevertheless, this unique experiment offers a detailed overview of needle structure and function across various environmental conditions.

Multi-year records of gas exchange at the individual tree-level in the same forest ecosystem are rare, and studies often focus either on short-term treatment effects or steady-state natural conditions. Nevertheless, long-term measurements are indispensable to distinguish intraspecific differences in photosynthesis capacity and sensitivity (Bachofen et al., 2020) and much of the uncertainty in projecting future terrestrial carbon uptake and storage is due to a lack of knowledge of the long-term response of photosynthetic carbon assimilation to future conditions (Friedlingstein *et al.*, 2014). With this long-term irrigation manipulation experiment in a natural forest, we studied the sensitivity to short-term environmental changes depending on long-term acclimation to soil water availability. We found that long-term acclimation to increased soil VWC has increased C assimilation on the leaf-level, which in combination with higher leaf area caused an increased C assimilation on the whole tree-level. This larger crown does not seem to make the leaves more sensitive to changes in atmospheric demand. Instead, drought release reduced the sensitivity of stomata to increasing VPD. Lastly, understanding how structural and biochemical adjustments occur due to environmental changes over time is indispensable for future predictions of how forests react to a changing climate. Thus, our findings that structural adjustments lead to an attenuation of initially strong leaf-level acclimation to strong multiple-year drought shed a new and important light on the memory effects and acclimation potential of evergreen trees to sudden environmental changes. The acclimation pathways found in this study are limited to a single pine species but are expected to be valid to a range of evergreen conifers, while research on deciduous species would greatly enhance our knowledge on the acclimation potential of forests all around the world.

## Acknowledgements

This study is based on data from the long-term Pfynwald irrigation experiment, which is part of the Swiss Long-term Forest Ecosystem Research Program LWF (www.lwf.ch). We thank Christian Hug, Peter Bleuler, Simpal Kumar, Peter Jakob, Matthias Häni, and Flurin Sutter for their continuing support on-site, for data preparation, and the fruitful discussions. A.G. and C.G acknowledge support from the Swiss National Science Foundation SNF (310030_189109 and PZ00P3_174068). Y.S. acknowledges support from NERC (RA0929) and the Finnish Academy (323843). Furthermore, we acknowledge the long-term help by Konrad Egger and his team from the Forstrevier Leuk, the Burgerschaft Leuk, and the HYDRO Exploitation SA.

## Author contributions

A.R., A.G., M.S., Y.S., and L.S. designed the experiment. M.S., Y.S., L.S., J.G. and P.D. carried out gas exchange measurements, K.M. and R.Z. were responsible for continuous soil and meteorological data acquisition and cleaning. L.S., C.G. and B.S. analyzed the data and L.S. wrote the manuscript. All authors contributed to the interpretation and writing of the manuscript.

## Data availability

The data that support the findings of this study are available from the corresponding author upon reasonable request.

