## Supporting information for "Photosynthetic acclimation and sensitivity to short- and long-term environmental changes"

**Fig. S1**. Vapor pressure deficit (VPD) outside *vs.* inside the cuvette. The diagonal dotted line indicates the 1:1 ratio.

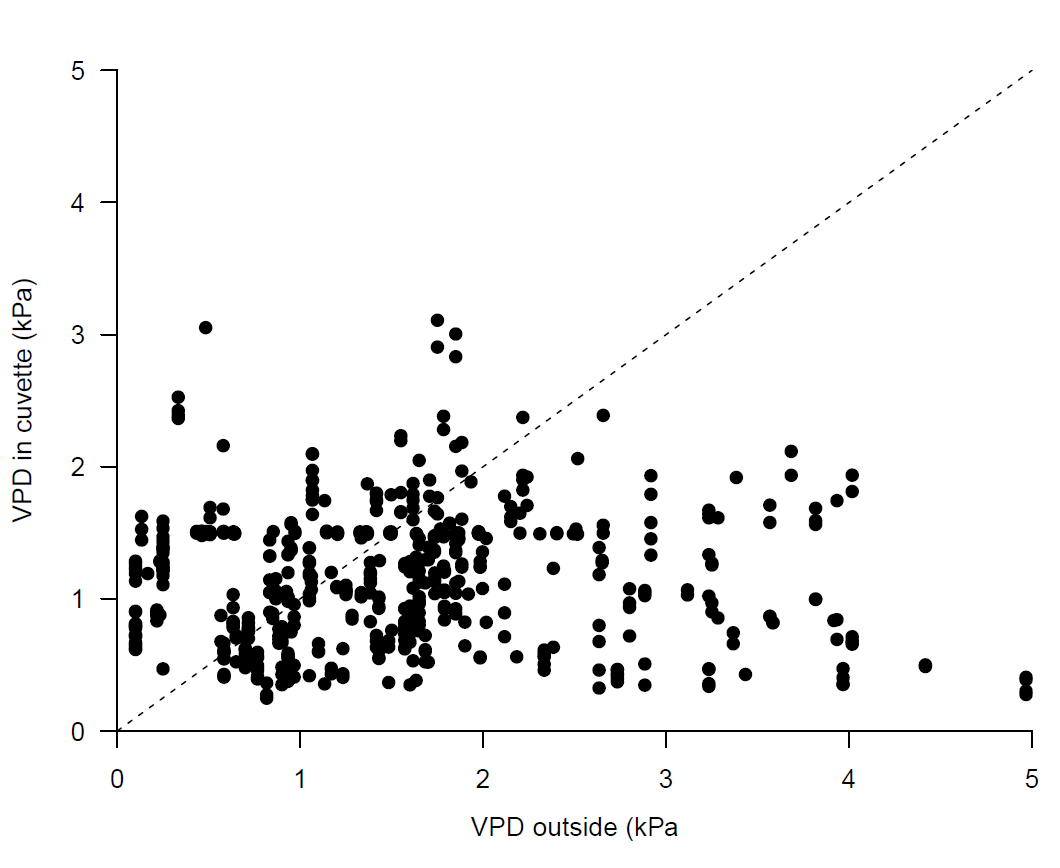

**Fig. S2.** a) Sensitivity parameter m of g_s_ to VPD in the three VWC bins as derived from the linear mixed effect model. b) Rasterplot of modeled differences between g_s_ of irrigated *vs.* control trees for a continuum of soil VWC and VPD conditions.

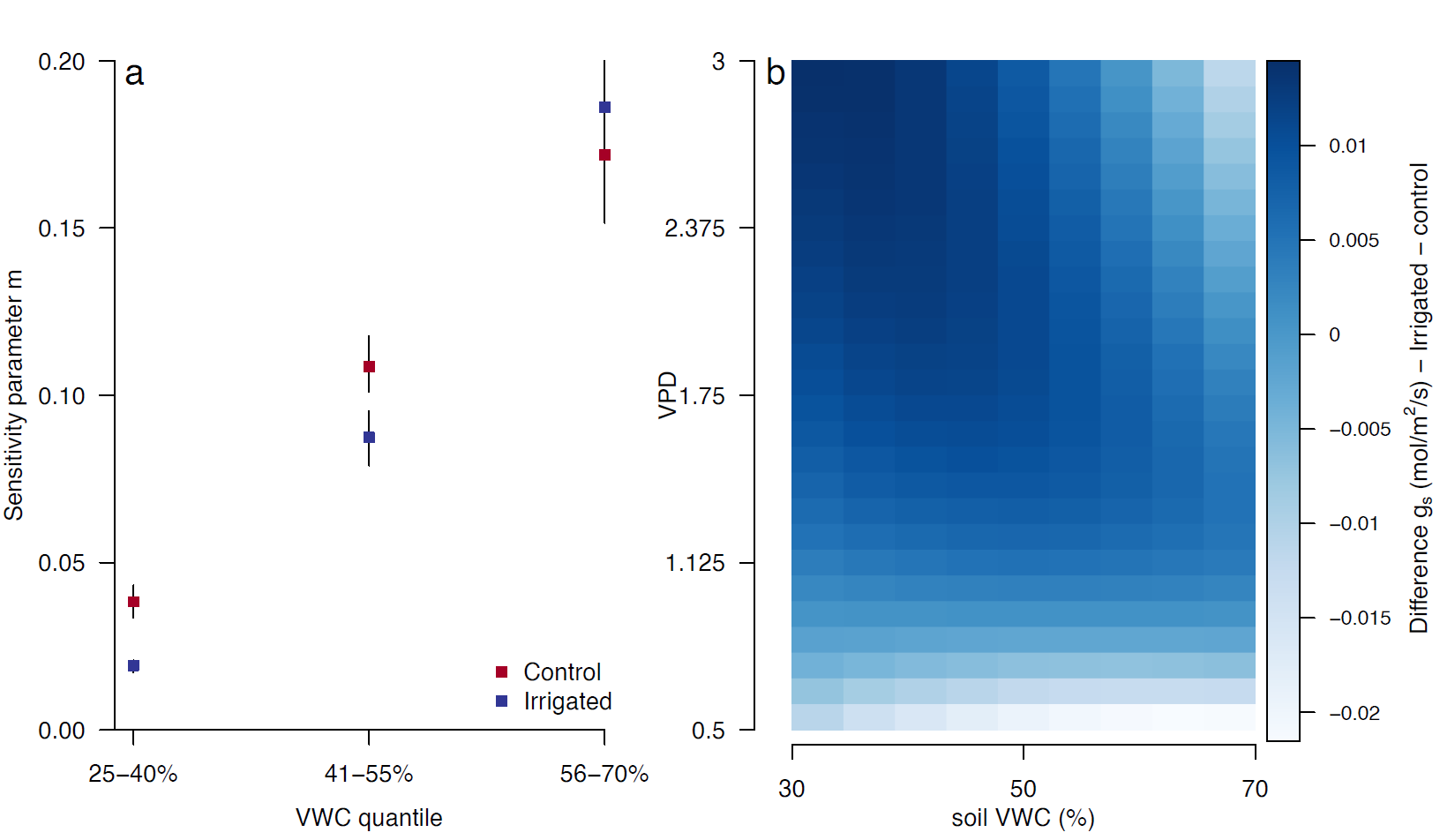

**Fig. S3.** Photosynthesis (A), stomatal conductance (g_s_), transpiration (E), intrinsic water use efficiency (WUE_i_), Vc_max_ and J_max_ and the ratio between the two in irrigated (blue dots) and irrigation-stop (orange dots) trees on all measuring dates between 2014 and 2019, during the time irrigation had stopped. Mean and SE are shown, and asterisks show significant treatment differences according to linear mixed effect models (p < 0.05).

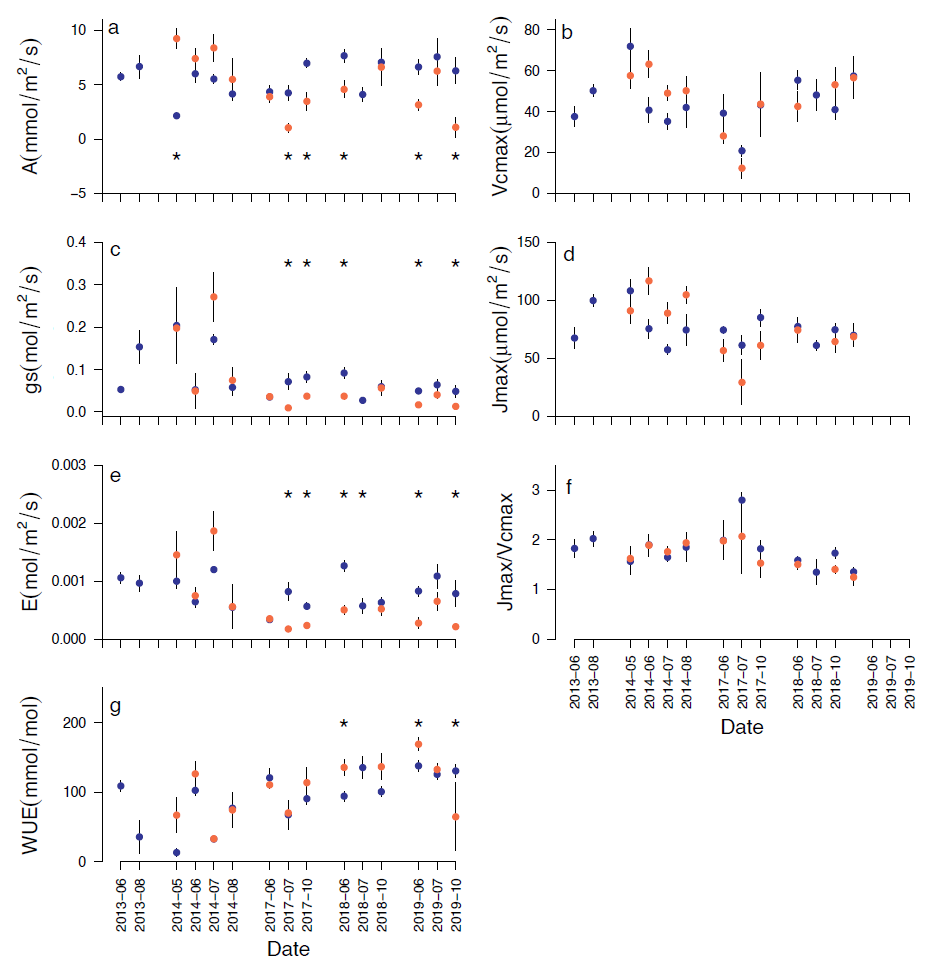

**Table S1.** Fitted parameters of Tp (Transition point), Rd (day respiration), Γ* (photorespiratory compensation point), and Km (Michaelis-Menten Coefficient) per treatment (Control, Irrigated, Irrigation-stop). Mean and SE are shown. Different letters in superscript indicate significant differences between treatments (p < 0.05).

|  | Tp | Rd | Γ* | Km |
| --- | --- | --- | --- | --- |
| Control | 455.11 ± 33.67^a^ | 0.49 ± 0.11^a^ | 35.49 ± 0.70^b^ | 531.02 ± 18.05^b^ |
| Irrigated | 492.81 ± 32.12^a^ | 0.71 ± 0.08^a^ | 38.79 ± 0.73^a^ | 615.69 ± 20.45^a^ |
| Irrigation-stop | 492.65 ± 53.51^a^ | 0.91 ± 0.14^a^ | 36.30 ± 0.87^ab^ | 549.30 ± 20.51^ab^ |

**Table S2**: Mean and standard error of gas exchange parameters in control (naturally drought exposed), irrigated and irrigation-stop trees. A (photosynthesis, mmol/m^2^/s), g_s_ (stomatal conductance, mol/m^2^/s), E (transpiration, mol/m^2^/s), WUE (water use efficiency, mmol/mol), Vc_max_ (Rubisco carboxylation, *µ*mol/m^2^/s), J_max_ (electron transport rate, *µ*mol/m^2^/s), J_max_/Vc_max_ (unitless). Different letters indicate significant differences between groups as tested by a mixed effect model with tree number as a random factor (p < 0.05).

|  | **Control** | **Irrigated** | **Irrigation-stop** | | | |
| --- | --- | --- | --- | --- | --- | --- |
|  |  |  | *2014* | *2017* | *2018* | *2019* |
| **A** | 4.69 ± 0.26^b^ | 6.33 ± 0.24^a^ | 7.50 ± 0.65^a^ | 3.17 ± 0.41^b^ | 5.24 ± 0.80^ab^ | 3.90 ± 0.54^b^ |
| **g_s_** | 0.06 ± 0.004^b^ | 0.08 ± 0.00^a^ | 0.14 ± 0.03^a^ | 0.03 ± 0.00^b^ | 0.04 ± 0.01^b^ | 0.03 ± 0.01^b^ |
| **E** | 0.51 ± 0.03^b^ | 0.79 ± 0.04^a^ | 1.15 ± 0.15^a^ | 0.28 ± 0.03^b^ | 0.49 ± 0.06^b^ | 0.47 ± 0.08^b^ |
| **WUE** | 111.64 ± 4.35^a^ | 104.92 ± 3.51^a^ | 88.72 ± 12.56^c^ | 100.23 ± 7.31^bc^ | 131.92 ± 9.37^ab^ | 147.89 ± 9.81^a^ |
| **Vcmax** | 44.38 ± 2.14^a^ | 43.88 ± 1.71^a^ | 56.58 ± 3.26^a^ | 26.64 ± 4.32^b^ | 51.52 ± 5.19^a^ | 56.56 ± 10.34^a^ |
| **Jmax** | 69.45 ± 2.94^a^ | 75.78 ± 2.47^a^ | 99.81 ± 5.42^a^ | 50.23 ± 6.91^b^ | 73.76 ± 6.67^ab^ | 68.61 ± 3.57^ab^ |
| **J/Vc** | 1.64 ± 0.05^b^ | 1.83 ± 0.06^a^ | 1.81 ± 0.09^a^ | 1.83 ± 0.16^a^ | 1.45 ± 0.06^a^ | 1.24 ± 0.16^a^ |

**Table S3**: Anova results of the best fitting linear mixed effect model (based on lowest AIC) for stomatal conductance (g_s_). The full model consisted of treatment, a 2^nd^ degree polynomial of soil volumetric water content (VWC) at 10 cm and the logarithm of vapor pressure deficit (VPD) in the cuvette, after which non-significant / non-contributing factors were discarded.

| **g_s_ (sqrt)** | **F** | **p** |
| --- | --- | --- |
| Treatment | 0.20 | 0.657 |
| Log(VPD) | **14.71** | **< 0.001** |
| VWC^2^ | **64.89** | **< 0.001** |
| VWC | **71.09** | **< 0.001** |
| Treatment:log(VPD) | **5.18** | **0.023** |
| VPD:VWC | **52.91** | **<0.001** |

**Table S4**: Coefficients for the best fitting linear mixed effect model (as shown in table S3). Values in bold indicate significance (p < 0.05) between treatments.

| **g_s_ (sqrt)** | Coefficient (SE) |
| --- | --- |
| Control | 1.747 (0.207) * VWC – 0.184 (0.047) |
| Irrigated | 1.747 (0.207) * VWC – 0.178 (0.014) |
| Control m (log(VPD)) | -0.629 (0.086) * VWC + **0.113** |
| Irrigated m (log(VPD)) | -0.629 (0.086) * VWC + **0.156** |
